# Deep learning-enhanced light-field imaging with continuous validation

**DOI:** 10.1101/2020.07.30.228924

**Authors:** Nils Wagner, Fynn Beuttenmueller, Nils Norlin, Jakob Gierten, Juan Carlos Boffi, Joachim Wittbrodt, Martin Weigert, Lars Hufnagel, Robert Prevedel, Anna Kreshuk

**Author notes:** These authors contributed equally to this work (Nils Wagner, Fynn Beuttenmueller, Robert Prevedel, Anna Kreshuk). Correspondence should be addressed to R.P. and A.K.

## Abstract

Light-field microscopy (LFM) has emerged as a powerful tool for fast volumetric image acquisition in biology, but its effective throughput and widespread use has been hampered by a computationally demanding and artefact-prone image reconstruction process. Here, we present a novel framework consisting of a hybrid light-field light-sheet microscope and deep learning-based volume reconstruction, where single light-sheet acquisitions continuously serve as training data and validation for the convolutional neural network reconstructing the LFM volume. Our network delivers high-quality reconstructions at video-rate throughput and we demonstrate the capabilities of our approach by imaging medaka heart dynamics and zebrafish neural activity.

---

Capturing neuronal activity distributed over whole brains, elucidating long-range molecular signaling networks or analyzing structure and function of beating hearts in small animals necessitate imaging methods that are capable of resolving these highly dynamic processes on milli-second time-, and hundreds of micrometer length scales. To address these challenges, several imaging approaches^1^ have recently been proposed or optimized, ranging from highly optimized point and line scanning, to selective plane illumination^2^ or by reducing the dimensionality of the image acquisition^3^. While the former two are limited by the sequential nature of the image capture, the latter often require sparsely labelled samples or a severely compromised imaging field-of-view (FOV).

A particularly attractive candidate for high-speed three-dimensional (3D) imaging in biology is light-field microscopy (LFM), due to its ability to instantaneously capture 3D spatial information in a single camera frame, thus permitting volumetric imaging limited by the frame-rate of the camera only^4–6^. The exceptional ability to image the 3D distribution of fluorescent emitters over large, hundreds of micrometer-scale FOV with millisecond temporal resolution has opened new avenues in developmental and neuro-biology, such as the recording of whole-brain neuronal activity in several model organisms^6–8^ or the visualization of ultra-fast cardiovascular dynamics^9,10^. Technologically, LFM has seen a steady raise in performance over the past years, including approaches to advance its rather low and non-uniform spatial resolution^9^ and signal-to-noise^10,11^, and to optically^9,12^ or computationally^13–15^ reduce the presence of image reconstruction artefacts. Yet the widespread use of this optically appealing technique in the life sciences has been hampered by a computationally demanding, iterative image reconstruction process that ideally demands large-scale computational infrastructure as well as data management, and thus practically restricts the effective experimental throughput, especially with respect to long-term recordings.

With the explosive development of deep learning and convolutional neural networks (CNNs), multiple algorithms have recently been proposed with the aim to replace iterative deconvolution procedures such as Richardson-Lucy’s by a CNN^16^. In the natural image domain, CNNs are now the primary method for removing motion blur and other artefacts traditionally solved by iterative deconvolution^16^. Similarly, in microscopy several deep learning-based methods have recently been introduced for deblurring, denoising or super-resolution applications^18^. Although these methods demonstrate excellent image reconstruction performance and empirically have been shown to generalize to data similar to the one used in training, no theoretical guarantees on generalization can be given. It is therefore of utmost importance to extensively validate and, if needed, to retrain the CNN for each experimental setting^19^. This requirement presents a problem for many bio-imaging applications, and in particular for dynamic imaging with LFM, as raw light-field images are difficult to interpret. For many dynamic biological processes, it is not possible to arrest the activity and acquire a static training volume by confocal or other imaging modalities^20,21^. In that case, training has to be restricted to simulations or iterative algorithm reconstructions, without showing the CNN any independently acquired volumetric data which truly corresponds to the light-field images it aims to reconstruct.

To overcome this limitation, here we present a novel framework for fast and high-fidelity reconstructions of experimental light-field microscopy images, termed HyLFM. Our approach is based on reconstruction by a CNN enhanced by simultaneous acquisition of high-resolution image data for training and validation. Our neural network – which we term HyLFM-Net – is designed for light-field data processing and 3D image reconstruction (see **SI Fig. 1** and **SI Tab. 1** for a detailed architecture description). To avoid potential bias to the previously seen training data and to enable direct validation of the reconstructions from uninterpretable LFM images, we have included an additional, continuous validation mechanism into our LFM imaging setup, thereby achieving and ensuring high-fidelity, trustable reconstructions. Experimentally, this is realized by adding a simultaneous, selective-plane illumination microscopy (SPIM) modality into the LFM setup which continuously <u>scans through the volume and</u> produces high-resolution ground truth images of single planes for validation, training or refinement of the CNN. The training can thus be performed both on static sample volumes and dynamically from a single plane that sweeps through the volume during 3D image acquisition. Besides direct training from non-static samples, the latter approach allows retraining the network if <u>its reconstructions do not agree with the individual SPIM plane images taken during</u> continuous validation. We demonstrate the capabilities of our HyLFM system by imaging the dynamics of a hatchling (8 dpf) medaka *(Oryzias latipes)* beating heart across a 350×300×150 μm^3^ FOV at a volumetric speed of 40-100Hz, as well as calcium-evoked neural activity in 5dpf zebrafish *(Danio rerio)* larvae over 350×280×120 μm^3^ FOV at 10Hz. HyLFM-Net achieves superior image reconstruction quality (and spatial resolution) compared to traditional, iterative LFM deconvolution-based techniques^5,6^ and at a video-rate (18Hz) inference throughput (**SI Tab. 2**). <u>Furthermore, we demonstrate performance and online validation of the network trained on static volumes or dynamically acquired individual planes, with and without additional network fine-tuning on a part of the dynamically acquired timelapse</u>.

The design of the HyLFM imaging system is conceptually shown in **Fig. 1a** and based on an upright SPIM configuration with dual-illumination (**Online Methods and SI Fig. 2**). This allows to simultaneously or sequentially illuminate the entire sample volume for light-field and/or a single plane for SPIM recording. On the detection side, the objective (Olympus, 20x 0.5NA) collects the excited fluorescence which is split either via a 30/70 beamsplitter or based on wavelength into separate optical paths for SPIM and LFM imaging, respectively. A fast, galvanometric mirror in the illumination path, together with an electrical-tunable lens (ETL) in the SPIM detection path enables to arbitrarily reposition the SPIM excitation and detection planes in the sample volumes at high speed (15ms), in respect to the LFM imaging volume (**Fig. 1b**). An automated image processing pipeline^9^ ensures that both LFM and SPIM volumes are co-registered in a common reference volume/coordinate system with high precision, which is an important prerequisite for CNN training and validation. Our ability to simultaneously acquire both 2D and 3D training data is paramount to ensure high-fidelity and reliable CNN light-field reconstructions of arbitrary samples, including data never seen in previous training. Furthermore, this includes dynamic samples for which the process of interest cannot be arrested to acquire a static training volume at high resolution. This is an important advancement of our system compared to previous work on artificial intelligence enhanced LFM reconstructions^20–22^.

**Figure 1:**
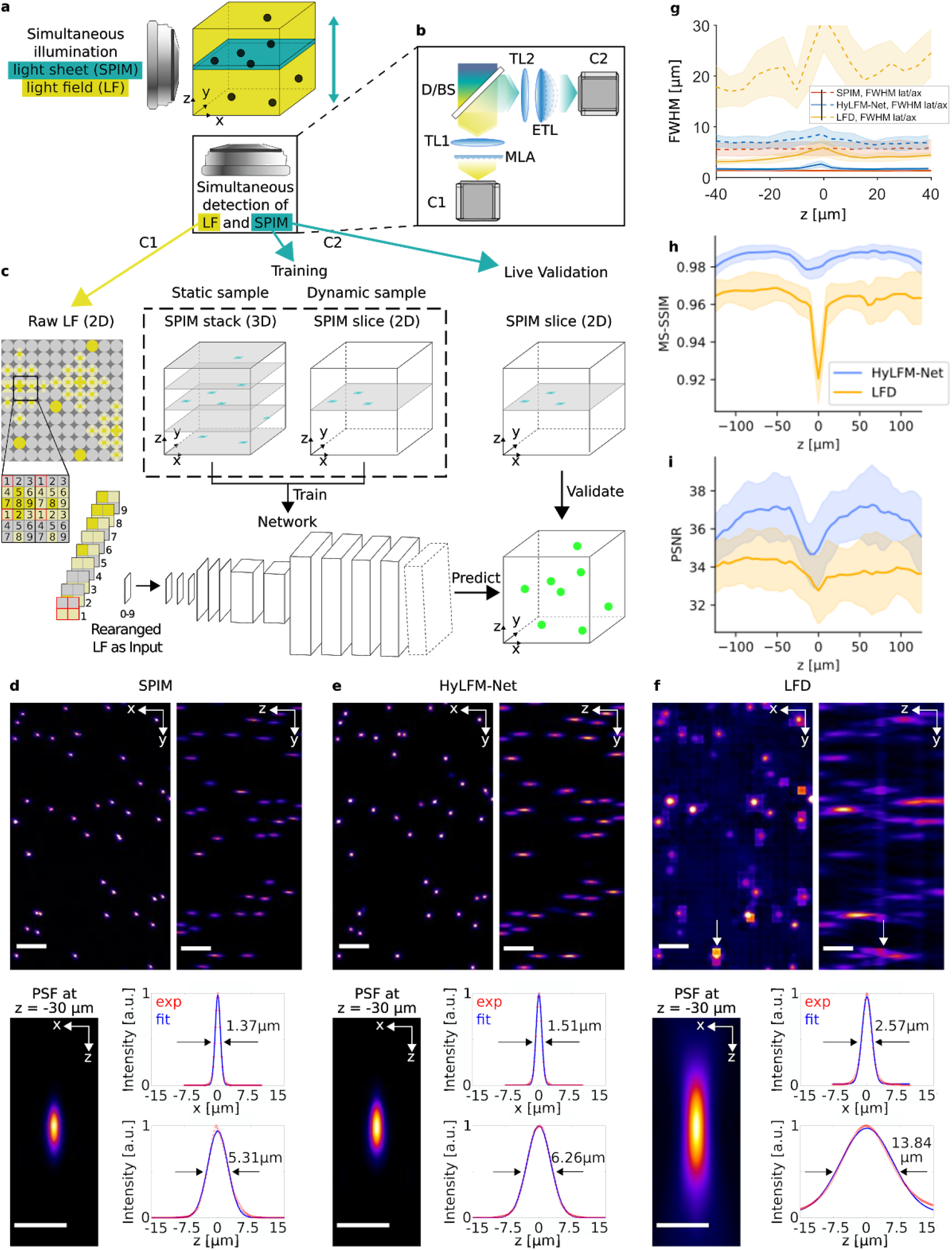
Schematic principle and experimental characterization of HyLFM imaging performance. **(a)** Microscope geometry showing simultaneous imaging via SPIM and light-field modalities. **(b)** Schematic view of the SPIM and light-field detection paths. (D-)BS, (dichroic) beamsplitter <u>used for (dual-) or single-color imaging</u>; TL, tube lens; MLA, microlens array; ETL, electrotunable lens; C, sCMOS camera. **(c)** HyLFM-Net image reconstruction pipeline. The raw light-field image serves as input to a reconstruction CNN, where the lenslet pixels are rearranged channelwise. The last layer encodes the affine transform between the SPIM and LFM spaces. The CNN can be trained either on high-resolution light sheet volumes, for static samples, or on high-resolution light sheet planes, for dynamic samples. The network output can additionally be validated by sweeping light-sheet planes. **(d–i)** Evaluation of HyLFM-Net’s performance on sub-diffraction sized, fluorescent beads. A 3D SPIM stack **(d)** and the corresponding raw light-field image are recorded. **(e)** Same volume is reconstructed by a HyLFM-Net trained on other SPIM / light-field pairs. **(f)** Conventional reconstruction based on Richardson-Lucy type light-field deconvolution (LFD). Exemplary artefacts are highlighted by white arrows. **(g)** Lateral and axial resolution as a function of imaging depth for SPIM, HyLFM-Net and LFD, respectively. MS-SSIM **(h)** and PSNR **(i)** image quality metrics across imaging volume comparing HyLFM-Net and LFD with the SPIM ground truth. <u>Shadows in JΛÜ. denote standard deviation, inferred from all beads in the dataset at the same z-position</u>. Scale bars in **(d-f)** are 20μm in whole FOV (top row) and 10μm in PSF close-up (bottom row). <u>PSF close-ups are computed by averaging over 10 beads. Experimental data in **(d-f)** is shown in red and fitting curves in blue</u>.

Deep learning-based image reconstruction methods are commonly trained from “original”-”reconstruction” image pairs, although semi-supervised and self-supervised approaches are now also gaining popularity^23,24^. Here, we follow a fully supervised approach and train HyLFM-Net directly on pairs of SPIM-LFM images. The LFM image which serves as input to the network is transformed into a tensor where the individual pixels of each lenslet are rearranged as channels (**Fig. 1c**). This re-arrangement <u>puts different angular views in different channels</u>. The convolution operations <u>can then</u> act on angular views, while the projection (1×1 convolution) layers learn to combine information from different angles. The multi-channel 2D images are passed through 2D residual blocks and a transposed convolution. The output goes through a final 2D convolution layer and is then transformed to 3D by reinterpreting network filters as the axial spatial dimension. The 3D images are then further processed by 3D residual blocks and upsampled by transposed convolutions to finally yield the reconstructed 3D volume. For training on single planes, the registration transform between the two detection modalities is encoded into the last network layer to enable direct comparison with the acquired 2D light sheet image (**Online Methods, SI Fig. 1**).

To evaluate and verify the performance of our HyLFM system, we imaged sub-diffraction sized, fluorescent beads suspended in agarose and quantified the improvement in both spatial resolution and overall image quality by comparing it with the standard, iterative light-field deconvolution (LFD – **Fig. 1d–i**) <u>as well as with the same deconvolution improved by a deep learning-based image restoration (Content Aware REstoration) method— (LFD+CARE – **SI Fig. 4**</u>). We found that HyLFM-Net correctly inferred the 3D imaging volume from the raw light-field data, yielding high and uniform lateral (1.8±0.2μm) as well as axial (7.1±1.3μm) resolution across the imaging volume (n=4966 beads, **Fig. 1d**), significantly better than what could be obtained by LFD (**Fig. 1i**). Furthermore, HyLFM-Net reconstructions do not suffer from artefacts near the native focal plane that are common in LFD^5^ (see arrows in **Fig. 1i** and **SI Fig. 3**). <u>While HyLFM-Net and LFD+CARE yield qualitatively comparable quality, the latter shares the same limitations of slow reconstruction speed as LFD alone. Furthermore, HyLFM-Net outperforms LFD+CARE in terms of object-level precision/recall metrics (**SI Fig. 5**</u>).

With all CNN based reconstruction methods, it has to be noted that the “PSF’’ of the signal reconstructed with a CNN depends strongly on the resolution and the shape of the signal in the training data. Therefore, training on small structures such as sub-diffraction sized beads will lead to unnaturally precise signal localizations and, when such a network is applied to a dataset with larger structures (or beads), can lead to erroneous reconstructions. Conversely, training only on large structures can lead to a network which merges small neighboring objects together (**SI Fig. 6**). While empirically the bias to training data can be alleviated by ensuring more diverse training datasets, at the current state of machine learning theory no formal guarantees on network generalization performance can be made. This observation has motivated our hybrid microscope setup, where the network can always be validated on concomitantly acquired high-resolution data and, if necessary, retrained directly on the imaged sample instead of relying on a sufficiently broad and general composition of static training data, which in practice is typically lacking or time- and/or resource-intensive to produce.

Next, we applied the HyLFM system to the challenging task of imaging a beating medaka fish heart *in-vivo* to show its capability to correctly capture highly dynamic cellular movements in 3D (**Fig. 2a–h** and **SI Video 1**). <u>Here we used dual color labelling of</u> cardiomyocytes *(myl7::H2B-eGFP; myl7::H2A-mCherry;* i.e. nuclear-eGFP for LFM-, -mCherry for SPIM detection-path) <u>to image the same features in SPIM and LFM with 40-100Hz volume rate. Doing so allowed us to</u> visualize the heart at single-cell resolution and free of reconstruction artefacts for both pharmacologically arrested (static) hearts (**Fig. 2a–c,** V-FOV ~350×300×180μm^3^), and beating (dynamic) hearts (**Fig. 2d–i,** V-FOV ~350×300×150μm^3^). Here, HyLFM-Net yielded high image quality metrics (multi-scale structural similarity index measure (MS-SSIM) = 0.87<u>1±0.007</u>, peak signal-to-noise ratio (PSNR) = 31.<u>6±0.6</u>) with regards to SPIM (**Fig. 2j-k**), while the comparison to LFD and image restoration setup again showed similar performance (**SI Fig. 7**). Importantly, note that the network trained on dynamically acquired SPIM single planes (HyLFM-Net-dyn in **Fig. 2j-k**) is performing equally well as the network trained on full static volumes (HyLFM-Net-stat in **Fig. 2j-k**). This underscores the feasibility of our hybrid 2D/3D imaging approach.

**Figure 2:**
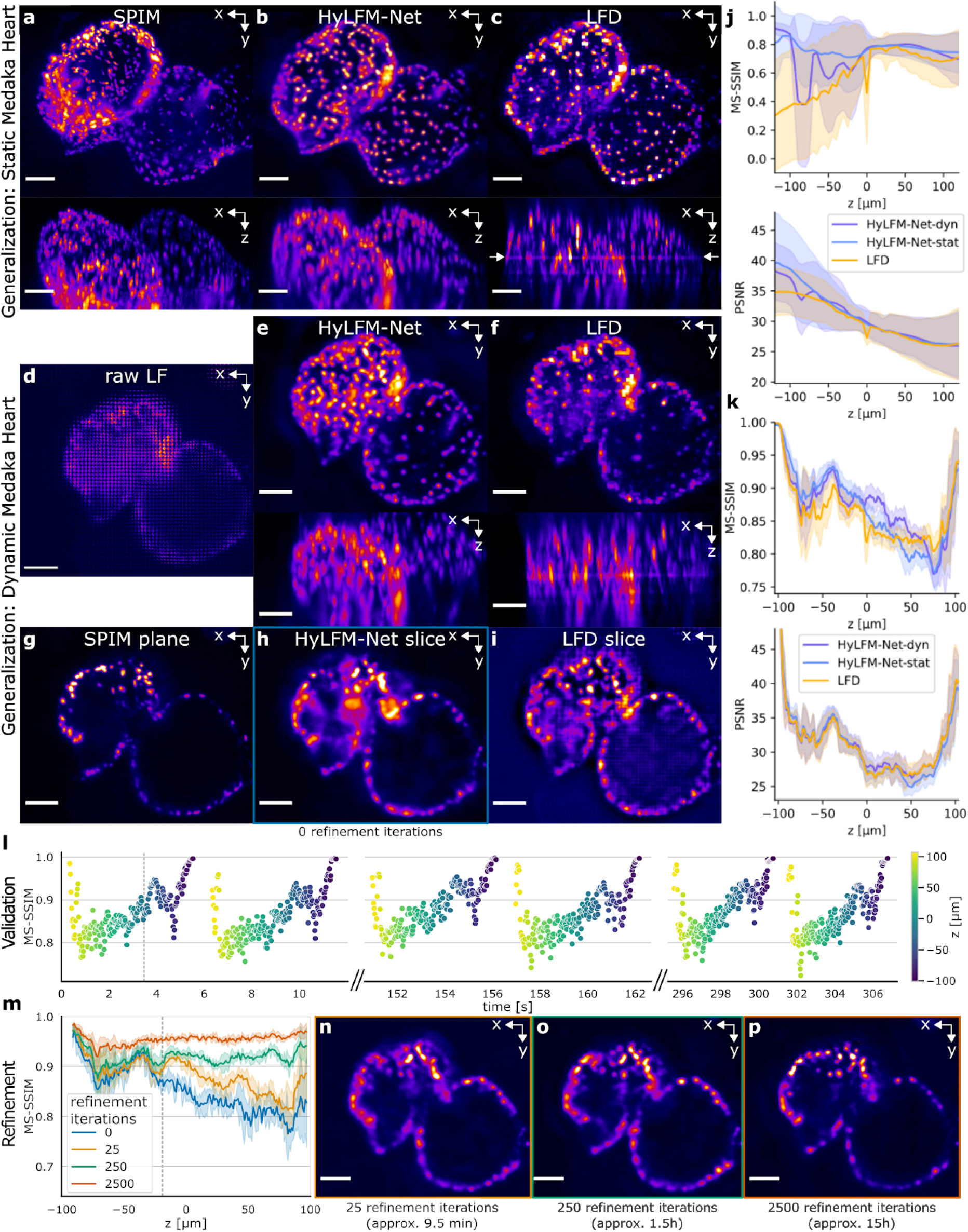
Experimental demonstration of HyLFM on medaka heart. **(a)** A static hatchling medaka heart acquired by light sheet (SPIM, maximum intensity projections). The corresponding light-field volume reconstructions by **(b)** HyLFM-Net and **(c)** LFD. <u>Here, HyLFM-Net (later HyLFM-Net-stat) was trained on different static volumes</u>. Note reconstruction artefacts (white arrows) and signal dimming in off-center regions in LFD. **(d)** Example raw light-field (LF) image of a dynamic (beating) medaka heart, acquired at 40Hz. **(e)** Maximum intensity projection of the volumetric HyLFM-Net-stat reconstruction <u>of the light-field image from **(d)**</u>. <u>Note improved cellular details and absence of artefacts compared to LFD reconstructions in **(f)**. **(g)** High resolution light sheet image plane at −19μm depth and same time-point as in **(d)**, used for single plane validation. **(h)** Slice of HyLFM-Net reconstruction corresponding to **(g)**. **(i)** Slice of LFD reconstruction corresponding to **(g)**. **(j-k)** HyLFM-Net-stat compared to HyLFM-Net trained on dynamically acquired individual planes (HyLFM-Net-dyn) and LFD. **(j)** MS-SSIM and PSNR image quality metrics across the imaging volume, compared to SPIM ground truth from **(a)**. **(k)** image quality metrics for reconstructions of the light-field image in **(d)**, compared to dynamic SPIM ground truth from **(g). (l)** Continuous validation plotting MS-SSIM of pre-trained HyLFM-Net-stat over imaging time to showcase data of **(k)** in an experiment-like setting. Color denotes axial position in volume. **(m)**_ MS-SSIM image quality metrics for increasing refinement iterations also visualised in **_(ņ ßL_** Note the obvious improvement of the reconstruction fidelity between pre-trained network without fine-tuning shown in **(h)** and fine-tuned reconstructions of **(n-p)**, Grey dashed lines in **(l,m)** indicate the timepoint/axial position shown in **(g)**, for which the three increasing refinements **(n–p)** are visualised. Shadows in **(j,k,m)** denote standard deviation, inferred from static test volumes **(j)** and from all test timepoints/single planes of a dynamic dataset **(k,m)**. Scale bar is 50μm in **(a–p)**. See also **SI Videos 1-3** for **(d-i)** and **SI Fig. 7** for a comparison to LFD improved by image restoration, as well as **SI Video 4** and **SI Figure 8** for visualization and more details on fine-tuning experiments</u>.

HyLFM-Net allowed 3D volume inference at 18.2 Hz, which represents at least a 1000-fold reconstruction speed improvement over common LFD^5,6^ (**SI Tab. 2**). <u>Here we highlight that all reconstructions and image quality metrics of **Fig. 2** were obtained in inference mode from networks trained on separate fish hearts, illustrating the high generalization ability of our HyLFM-Net</u>. Being able to perform 3D volume inference at video-rate speed significantly boosts overall experimental imaging throughput and further enables real-time ‘quality’ control of uninterpretable light-field images during acquisition. <u>To demonstrate this aspect of on-the-fly validation, we compute MS-SSIM of the volume slice corresponding to the simultaneously acquired scanning SPIM plane (**Fig. 2l**). This allows us to continuously monitor – in real time – the image quality of our network reconstruction and to check whether the metrics fall short of a user-defined threshold that depends on the reconstruction accuracy requirements of the experiment (e.g. MS-SSIM<0.8). In such a case, network fine-tuning can be performed based on the single-plane SPIM images as training data (**Fig. 2m-p, SI Fig. 8**). To illustrate this fine-tuning process on dynamically beating medaka heart data, we started with a network which was trained on a static heart. Then we used 8883 individual SPIM planes from a 5-minute timelapse as training data and further 756 unseen consecutive planes from the same timelapse to evaluate the performance. After just 25 iterations (10 minutes) of re-training image quality improved substantially (**Fig. 2m,n**), reaching optimum metrics after 2500 iterations (15 hours of GPU time, **Fig. 2m,p**) Importantly, we note that in principle no ongoing imaging experiment needs to be stopped for fine-tuning, as this can be performed on the already acquired imaging data. For longer fine-tuning runs which start from a network pre-trained on very different data (**SI Fig. 8**), the network can be updated during post-processing and therefore still provide high-quality reconstruction for analysis after the actual image acquisition has finished, while future experiments of the same kind will likely only need a few minutes of fine-tuning</u>.

The unique assets of LFM make it a promising method for neural activity imaging in small model organisms. To demonstrate the potential of HyLFM to also deliver quantitatively accurate reconstructions, we imaged 5dpf transgenic larval zebrafish brains expressing the nuclear-confined calcium-indicator GCaMP6s *Tg(elavl3:H2b-GCaMP6s)* (**SI Fig. 9**). When distributing the excited fluorescence into the LFM and SPIM detection arms we could record whole-volume light-field and high-resolution SPIM imaging data at 10 Hz each, over a 350×280×120μm^3^ volume. Again, the concurrent availability of ground truth data enabled our HyLFM system to faithfully learn and infer not only structural, but also intensity-based information, as demonstrated by the high degree of correlation of Ca^2+^-signal traces between HyLFM and ground truth data obtained by SPIM or conventional LFD (**SI Fig. 9b,c**). The ability to rapidly acquire neural activity dynamics from low signal light-field data at hundreds of Hz volume rate should make HyLFM an attractive method for visualizing electrical activity via recently developed voltage-indicators^25^, in which kHz-rate volumetric image data needs to be efficiently captured and reconstructed.

In summary, we have demonstrated a new framework for deep-learning based microscopy with continuous ground-truth generation for enhanced reconstruction reliability. Our approach enables light-field imaging with improved spatial resolution and minimal reconstruction artefacts, and compared to previous work based on multiview deconvolution^9^, achieves this performance within the relaxed imaging geometry of a standard two-objective SPIM. <u>For simultaneous SPIM and LFM imaging such as shown in Fig. 2, the sample requires dual-color labelling so that the modalities can be split by wavelengths (Fig. 1a). This setup is only limited in speed by the signal and/or frame rate of the camera readout. Alternatively, for single-color imaging the modalities need to be rapidly alternated and the fluorescence split stochastically. In this setup, the signal intensity in each modality is reduced and a slight time delay is introduced between acquisitions, depending on camera speed and fluorescence signal strength (e.g. in our case SPIM and LFM acquisitions are at least 27ms apart, see Methods). This could in principle lead to ambiguous network performance and validations for highly dynamic samples. Although the initial training time of HyLFM-Net from scratch can be long, our fine-tuning experiments suggest an approach to avoid it becoming a bottleneck: HyLFM-Net trained on beads can be converted to reconstruct dynamic biological samples in 10 hours of training on a single GPU. After retraining, the network can deliver optimal results for the target system and – in case of a change in the imaging conditions – can be retrained again on a new sample in mere minutes</u>.

The ability to reconstruct light-field volumes at sub-second (video-)rate eliminates the main computational hurdle for light-field imaging in biology, and we thus expect this to further accelerate the uptake of LFM by the community. The integration of a high-resolution imaging modality into our LFM system further mitigates the omnipresent problem of acquiring appropriate training data, as it can be generated simultaneously and on-the-fly. Furthermore, the on-line availability of single-plane ground truth (SPIM) images distributed over 3D space and time enables continuous CNN output validation and fine-tuning, as early time points of a time-lapse imaging experiment can be used for network training and/or refinement. This new concept to supervised AI-enhanced microscopy also solves the problem of transferability, as the network over time learns on the actual experimental data, and therefore does not require pre-acquisition of training images from particular specimen types with separate microscopes. This is a key advantage of our approach which goes beyond previous work in the field^20–22^. While the HyLFM setup has been developed specifically for light-field imaging, the general principle behind it is applicable to other imaging approaches which rely on iterative or trained computational methods for image reconstruction or restoration. Finally, we note that access to time- and resource-efficient light-field reconstructions further facilitates data-intensive, long-term 4D imaging experiments at high throughput as it allows to store volumetric image data in compressed form, i.e. as raw 2D light-field images^20^. Given that light-field detection only requires the moderately complex and inexpensive addition of a suitable microlens array into the imaging path and is in principle compatible with any custom or commercial SPIM realization, bears further potential for widespread use of this method in the life sciences, <u>especially in the context of high-repeats and -throughput imaging studies</u>.

## ACKNOWLEDGMENTS

We would like to thank the EMBL Heidelberg mechanical and electronic workshop for help as well as the IT Services Department for HPC cluster support and Christian Tischer from CBA for his help with volume registration. We further thank M. Majewsky, E. Leist and A. Saraceno for fish husbandry. We thank Krasimir Slanchev and Herwig Baier (MPI Martinsried) as well as Maximillian Hoffmann and Benjamin Judkewitz (Charite Berlin) for providing calcium reporter zebrafish lines. J.G. was supported by a Research Center for Molecular Medicine (HRCMM) Career Development Fellowship (CDF), the MD/PhD program of the Medical Faculty Heidelberg, the Deutsche Herzstiftung e.V. (S/02/17), and by an Add-On Fellowship for Interdisciplinary Science of the Joachim Herz Stiftung and is grateful to M. Gorenflo for supervision and guidance. N.N acknowledges support from Åke Wiberg foundation, Ingabritt and Arne Lundberg foundation, and Sten K Johnson foundation. This work was supported by the European Molecular Biology Laboratory (F.B, N.W., N.N., L.H, A.K., R.P.).

## AUTHOR CONTRIBUTIONS

A.K., L.H. and R.P. conceived the project. N.W. and N.N. built the imaging system and performed experiments with the help of J.G. J.G. generated transgenic animals under guidance of J.W. F.B., A.K. and M.W. conceived the CNN architecture. F.B. and N.W. implemented the CNN and other image processing parts of the computational pipeline and evaluated its performance. <u>J.C.B. performed Ca^2+^-data analysis</u>. A.K. and R.P. led the project and wrote the paper with input from all authors.

## Online methods

### Hybrid LFM-SPIM imaging setup

The microscope consists of one illumination and one detection objective, orthogonal to each other (see **SI Fig. 2**). The illumination sources are continuous-wave lasers (λ = 488 nm, 20mW, Omicron, and λ = 561 nm, 50 mW, Cobolt). We use a 10×0.3 NA (Nikon CFI Plan Fluor 10XW) water dipping objective for illumination and a 20×0.5 NA (Olympus UMPLFLN 20XW) water dipping objective for detection. For the latter, a tube lens with focal length of 200mm (Nikon MXA20696) yields an effective magnification of 22.5x. Two illumination paths, combined by a dichroic mirror (D1), enable simultaneous dual color light-sheet and light-field illumination. For single-color calcium imaging, two separate 488 nm excitation lasers were used and the dichroic mirror (D1) was replaced by a non-polarising beamsplitter (Thorlabs, 70:30). A digital light sheet was generated using one of the axes of a 2D galvo pair (Cambridge technology) combined with a scan lens (SL) and tube lens (TL1, 200mm). To achieve a selective volume illumination for light-field excitation, the laser beam was first expanded and the central plateau of the (Gaussian) illumination profile was used to illuminate the entire volumetric FOV at once. The lateral extensions of the volumetric FOV could be adapted by changing the size of an aperture (2D slit) that was used to crop out the central region of the laser beam. A lens (TL2, 300mm) focused the light on the objective back aperture. On the detection side, the microscope has two arms separated by either a dichroic mirror (D2) for dual color imaging or a non-polarising beamsplitter (Thorlabs, 2”, 70:30 ratio) for single-color (e.g. calcium) imaging. For the light-field detection arm, light first passes through a chromatic filter (BP2), and thereafter through a microlens array (pitch 125 μm and focal length 3.125 mm, RPC photonics MLA s125 f25) mounted in a six-axis kinematic mount (Thorlabs, K6XS) allowing fine adjustment of the array in respect to the optical axis. The microlens array is subsequently imaged onto a 4.2 megapixel (2,048×2,048 pixels) sCMOS camera (Andor Zyla) using a 1:1 relay macro lens objective (Nikon AF-S 105 mm 2.8 G VR IF-ED Micro). The second detection arm is used for recording of the light sheet image modality. The light sheet images were acquired by displacing the illuminated light sheet with respect to the focal plane with a galvanometric mirror while refocusing the detection plane remotely using an electrically tunable lens (ETL, Optotune EL-10-30)^26^. Here, the fluorescence first passes through a band pass filter (BP1), and the primary image plane is then relayed through the remote focusing unit using a 100mm lens (RL1) to a lens pair consisting of a −75mm offset lens (OL), and the ETL combined with another 100mm lens (RL2). Finally, two 150mm lenses (RL3, RL4) in a 4f configuration as well as a 1:1 macro lens (Nikon AF-S 105 mm 2.8 G VR IF-ED Micro) relay the image plane onto a second sCMOS camera (Andor Zyla). Precise sample positioning was enabled by a composite xyz linear positioning stage (Newport M-562-XYZ) together with a piezo stage (Nanos LPS-30-30-1-V2_61-S-N and controller MC101) and a small rotation stage (Standa, 7R128). Further information on the setup can be found in **SI Fig. 2**.

### HyLFM image acquisition and registration

The training data for our HyLFM-Net network consists of SPIM data (ground truth) and the corresponding 2D light-field image (input). Two options can be pursued to acquire high-resolution training data in our HyLFM setup: 1) The SPIM imaging plane remains stationary in order to sample the dynamics (e.g. multiple heart beats) over time and obtain sufficient variability in training data (e.g. heart shapes during the beat cycle) <u>before if moves to the next axial position in the volume and repeats the data acquisition</u>. 2) The SPIM modality continuously loops in 3D in order to acquire enough variability at each respective z plane. The second option minimizes potential photobleaching and -toxicity effects and was thus chosen for the majority of the experiments. Due to experimental imperfections, the FOV of the two detection paths might not overlap completely. In order to register the light sheet data to the light-field volume, we acquired a light-field image of fluorescent beads and a light sheet stack of the same volume by displacing the light sheet with respect to the focal plane with a galvanometric mirror and refocusing on the illuminated plane with the ETL. The light-field volume was then reconstructed from the recorded light-field image using Richardson-Lucy deconvolution (LFD) as in Ref.^6^. The two corresponding volumes were registered with the Multiview Reconstruction Plugin in Fiji^27^, yielding the affine transformation that maps the light-field volume to the light sheet stack. In dynamic training this affine transformation is then used within the final layer of the network and only the slice for which a light sheet equivalent has been acquired is sampled from the predicted volume during training. This routine also allows for an easy comparison between SPIM images and volumes reconstructed by the HyLFM-Net and LFD. We used the affine transformation that was computed from the fluorescent bead sample throughout all experiments.

### HyLFM-Net for light-field reconstruction

The input to the network is a 3-dimensional tensor. The original 2D light-field images, composed of up to 70×85 lenslets, 19×19 pixels each, are rearranged to contain 361 (19^2^) channels, with each channel corresponding to an angular view, i.e. same pixels of each lenslet. The input is normalized by its 5.0th and 99.8th percentile without clipping. For training and evaluation, the light sheet target images are normalized by their 5.0th and 99.9th percentile. During training several data augmentations are applied. These include addition of Gaussian or Poisson noise, joined random rescaling of light-field input image and light sheet target image, lateral axis flipping, as well as joined random 90-degree rotations (applied before rearranging light-field to 361 channels). The full network architecture is shown in **SI Fig. 1** and **SI Tab. 1**. Briefly, the rearranged 3D input tensor is passed through two or three residual blocks interlayered with transposed convolutional layers scaling up by factor 2 in the lateral dimensions. The output of the last residual block undergoes a final 2-dimensional convolution, after which its channel dimension is re-interpreted as an axial dimension and a smaller channel dimension. This 4D tensor is passed through 3D residual blocks and transposed convolutional layers, further upsampling in the two lateral dimensions. For predictions aligned with the SPIM data, the last layer of the network encodes the registration of the reconstructed LF volume to the SPIM volume. For static volumes the SPIM volume was transformed instead. In dynamic training only the one slice, for which a light sheet equivalent has been acquired, is sampled from the predicted volume. The network is trained either with L2 Loss or with a weighted, smooth L1 Loss (down-weighting non-peak-signal-pixels with a decaying weight). Only for the sparse bead data this choice has a significant impact on network convergence, as with the L2 loss the network converges to only predict background. The Adam optimizer is used with the learning rate set between 1.0e-5 and 3.0e-4. Training times for the networks are: 26.5h for HyLFM-Net-beads (beads), 121.2h for static heart (stat), 48h for dynamic heart (dyn), and 89.8h for neural activity tasks (brain), which were determined by observing the smooth L1 validation loss (beads) or the MS-SSIM validation score (stat, dyn, brain). <u>Final MS-SSIM metrics on the respective training datasets are 0.953±0.003 (beads), 0.93±0.04 (stat), 0.909±0.009 (dyn), and 0.90±0.02 (brain). As illustrated in **Fig. 2**, a refined network based on HyLFM-Net-stat was additionally trained with a mini-batch size of 8 and smooth L1 Loss for up to 6 epochs on 8883 dynamic heart slices scanning the entire volume 47 times. The refined network was tested on 756 dynamic heart slices from the same imaging session. It improved the MS-SSIM on its training/test data (both unseen test data for the unrefined HyLFM-Net-stat) from 0.868±0.005/0.86±0.01 to 0.971±0.002/0.970±0.003.</u>

<u>Similarly, the network shown in **SI Fig. 9** (HyLFM-Net-brain) was refined on 100 planes at a fixed axial position for 30 epochs with a mini-batch size of 5 (ca. 1.3h training time), which improved the test data (500 planes) from 0.90±0.02 to 0.938±0.003. The refined predictions on the test planes were then used to evaluate Ca^2+^trace correlations (see section Calcium imaging analysis).</u>

All training was done on a single NVIDIA GeForce RTX 2080 Ti GPU. Training data volumes measured 137 261.2×400.2×100 μm^3^ stacks (beads), 111 339.0×394.6×245 μm^3^ and 34 311.3×483.6×245 μm^3^ (stat), 13747 339.0×394.6 μm^2^ and 5925 277.9×283.5 μm^2^ slices (dyn), as well as 24502 355.7×439.1 μm^2^ slices (brain) (see also training data acquisition).

### Reconstruction quality analysis

To quantify the microscope’s performance in terms of spatial resolution, we imaged a 3D distribution of 0.1 μm sized fluorescent beads (TetraSpeck, Thermo Fisher Scientific) embedded in agarose. The Fiji plugin ‘Multiview-Reconstruction’^27^ was used to detect 3D bead positions in the recorded light sheet stacks. The same positions were then used to fit a 3D Gaussian and to compare the full width at half maximum (FWHM) in SPIM, LFD and HyLFM-Net prediction volumes respectively. <u>The PSF close-ups in **Fig. 1d-f** were computed by averaging over the same 10 beads for each modality at z=-30μm, using the MOSAIC Fiji plugin</u>. In order to investigate bias to training data and shape priors we imaged 4 μm sized fluorescent beads (TetraSpeck, Thermo Fisher Scientific), cross-applied trained deep neural networks and computed FWHM for all possible combinations (see **SI Fig. 6**). We computed MS-SSIM and PSNR values per z plane for light-field and network predictions, using light sheet planes as the reference, for the fluorescent beads and the medaka heart respectively. The following values were used for MS-SSIM computations: NumScales=5, ScaleWeights=fspecial(‘gaussian’, [1, numScales], 1), Sigma=1.5 (Ref.^28^). All reconstructions were rescaled for minimal MSE compared to SPIM.

### Fish husbandry and transgenic lines

All medaka fish are maintained in closed stocks at Heidelberg University. Medaka *(Oryzias latipes)* husbandry (permit number 35-9185.64/BH Wittbrodt) and experiments (permit number 35-9185.81/G-145/15 Wittbrodt) were performed according to local animal welfare standards (Tierschutzgesetz §11, Abs. 1, Nr. 1) and by European Union animal welfare guidelines. The fish facility is under the supervision of the local representative of the animal welfare agency. Medaka was raised and maintained as described previously^29^. For in-vivo imaging, embryos were kept in 165 mg/l 1-phenyl-2-thiourea (PTU) in embryo rearing medium (ERM) from 1 dpf until imaging to inhibit pigmentation. For heart imaging, the following transgenic medaka lines were crossed: *myl7::H2B-eGFP* (see Ref.^9^) and *myl7::H2A-mCherry.* For the generation of the *myl7::H2A-mCherry* transgenic medaka line the *myl7::eGFP* cassette of the pDestTol2CG plasmid (http://tol2kit.genetics.utah.edu/index.php/PDestTol2CG) was replaced by a *myl7::H2A-mCherry* cassette and the modified plasmid was co-injected with Tol2 transposase mRNA into wild-type stock Cab embryos as described earlier^30^. The calcium imaging experiments were performed using a zebrafish *(Danio rerio)* line with a nuclear-localized calcium sensor *(Tg(elavl3:H2B-GCaMP6s)).*

### Medaka imaging

Medaka larvae were imaged 1-3 days after hatching. Hatchlings were anesthetized in 150 mg/l disodium phosphate-buffered (pH 7,3) tricaine and mounted in 1 % low-melting agarose (in ERM) containing 150 mg/l tricaine. For light-field volume reconstructions with Richardson-Lucy deconvolution, a light-field PSF was chosen that yielded 49 distinct *axial* planes, spaced 5 μm apart after 8 iterations of deconvolution.

To create a static medaka heart for HyLFM-Net training, *myl7::H2B-eGFP, myl7::H2A-mCherry* transgenic medaka hatchlings were sedated with 150 mg/l tricaine and the heart was pharmacologically arrested with 40 mM 2,3-butanedione 2-monoxime (BDM). Pre-treated hatchlings were mounted in 1% low-melting agarose (in ERM) containing 150 mg/l tricaine and 40 mM BDM. In the case of re-onset of cardiac contractions, BDM stock solution (100 mM) was titrated into the sample chamber until the heart stopped beating. Then light-field images and light sheet stacks were acquired subsequently for the same position of the static heart. In order to get sufficient variability for training data we imaged the heart at multiple positions and at different angles. For this purpose, a linear piezo stage was used, which displaced the static heart diagonally to the detection objective, thereby assuring variations in two coordinates simultaneously, while the sample angle was modified manually with a rotation stage. For imaging the beating heart, we simultaneously acquired pairs of light-field and light sheet images at all z-planes in the volume. <u>For the SPIM modality, we scanned through the entire volume with 241 individual planes (spaced 1μm apart). At an effective SPIM frame rate of 40Hz (15ms refocusing + 10ms exposure time) the entire heart was thus imaged every 6 seconds</u>.

### Zebrafish calcium imaging

Zebrafish calcium imaging was performed using a nuclear localised calcium reporter Tg(elavl3:H2b-GCaMP6s). The zebrafish embryos were mounted according to previous work^9^ in 1 % low-melting agarose and imaged 5 days after fertilization using alternatingly SPIM and light-field illumination with 10Hz for both modalities.

<u>Training, validation, and test data were recorded by alternatingly acquiring light-field and SPIM images at 10Hz each (15ms exposure time + 12ms read-out time for SPIM, followed by 61ms exposure + 12ms read-out time for LFD). To cover the whole volume the SPIM plane was swept along the axial dimension for training and validation data. As test data we recorded consecutive time points at a fixed axial position to capture Ca^2+^ —transients</u>.

### Calcium imaging analysis

<u>SPIM and light-field collected datasets were motion-corrected by piecewise rigid motion correction package NoRMCorre. The motion corrected SPIM dataset was used to segment ROIs using custom Fiji macros. To do this, a standard deviation projection was obtained every 10 frames and the maximum projection of these was used to semi-automatically segment ROIs by thresholding. Signal threshold was set such that the signal obtained from ROIs was at least 2 times the standard deviation of the whole FOV. This ROI set was used to extract the Ca^2+^signal from both the SPIM dataset and the corresponding light-field datasets. We calculated the z-score of extracted raw Ca^2+^signals and smoothed them using a 10-point moving average. Traces from ROIs with no clear Ca^2+^transients in the SPIM dataset were excluded from further analysis. The Pearson correlation coefficient between SPIM traces and their corresponding light-field traces was calculated and compared using the Dunn-Sidak test. Normality was verified using Kolmogorov-Smirnov test. p < 0.05 was considered significant</u>.

## Code availability

The neural network code with routines for training and inference are available at https://github.com/kreshuklab/hylfm-net.

## Data availability

The datasets generated and/or analysed during the current study will be made publicly available at the point of publication.

