## Supplementary figures for "Deep learning-enhanced light-field imaging with continuous validation"

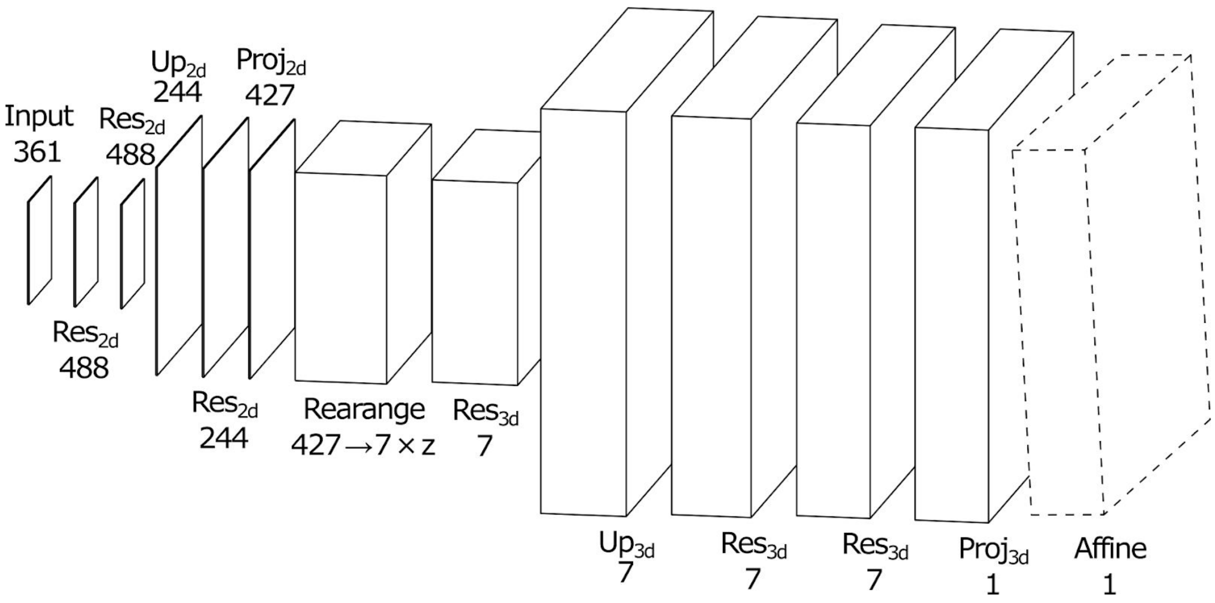


**Supplementary Figure 1: Network architecture.** $Res_{2/3d}$: residual blocks with 2d or 3d convolutions with kernel size (3×)3×3. Residual blocks contain an additional projection layer (1×1 or 1×1×1 convolution) if the number of input channels is different from the number of output channels. $Up_{2/3d}$: transposed convolution layers with kernel size (3×)2×2 and stride (1×)2×2. $Proj_{2d/3d}$: projection layers (1×1 or 1×1×1 convolutions). The numbers always correspond to the number of channels. With 19×19 pixel lenslets ($n_{num}$=19) the rearranged light field input image has $19^2$=361 channels. The affine transformation layer at the end is only part of the network when training on dynamic, single plane targets; otherwise, in inference mode it might be used in post-processing to yield a SPIM aligned prediction, or the inverse affine transformation is applied to the SPIM target for static samples to avoid unnecessary computations.

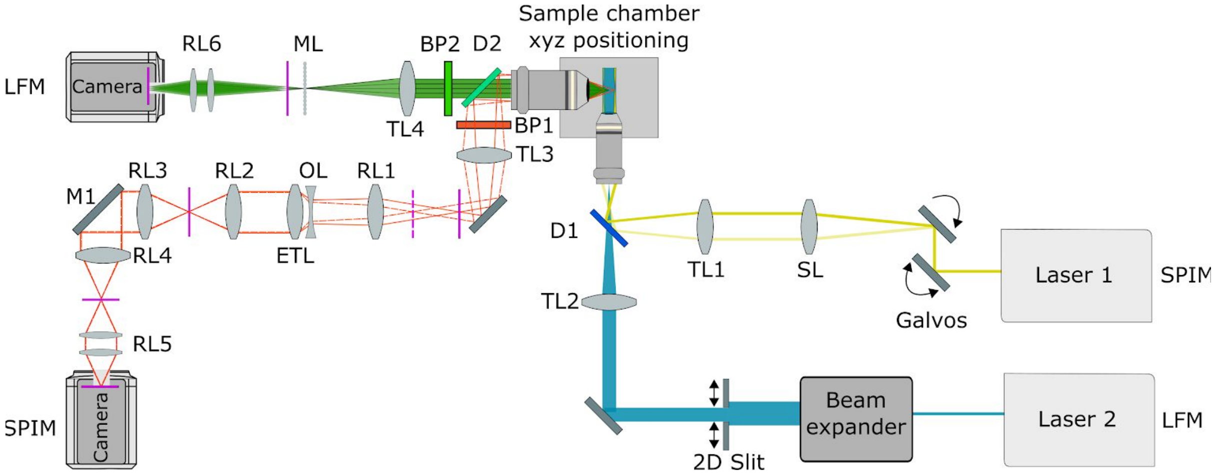


**Supplementary Figure 2: LFM-SPIM optical setup.** Schematic 2D drawing of the LFM-SPIM setup showing the main opto-mechanical components. The sample is illuminated through a single illumination objective with two excitation beam paths (ocra, light sheet illumination and blue, light field selective volume illumination) combined by a dichroic mirror (D1). The fluorescence is detected by an orthogonally oriented detection objective and optically separated onto two detection arms with a dichroic mirror (D2). Bandpass filters (BP1 and BP2) are placed in front of a tube lens (TL3,TL4) for the respective detection path. For the light field detection path (green), the tube lens (TL4) focuses on the microlens array (ML) and the image plane (shown in magenta) displaced by one microlens focal length is relayed by a 1-1 relay lens system (RL6) to an image plane coinciding with the camera sensor (shown in magenta).

For the light sheet detection path, a combination of several relay lenses (RL1 to RL4), a 1:1 macro lens (RL5) together with a lens pair consisting of an offset lens (OL) and an electrically tunable lens (ETL) is used to image two axially displaced objective focal planes (shown in magenta, dotted and solid) to a common image plane at the sensor. The refocusing is achieved by applying different currents on the ETL. The mirror M1 is placed at a Fourier plane, such that the FOV of the light sheet path can be laterally aligned to fit the light field detection FOV. For single color imaging, the dichroic mirrors D1 and D2 are replaced by beamsplitters. See Methods for details.

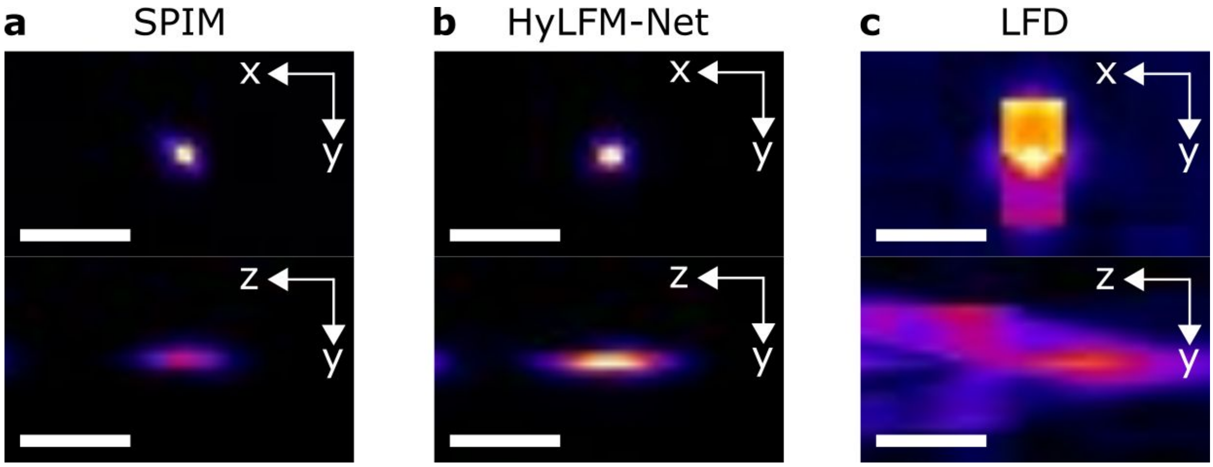


**Supplementary Figure 3: HyLFM-Net provides artifact free deconvolution of LFM data.**
**(a)** Ground truth single light sheet image. **(b)** Subdiffraction beads reconstructed by LFM-Net and **(c)** iterative light field deconvolution (LFD). The volume reconstructed by LFD shows clear signs of artifacts which are removed in HyLFM-Net reconstructed volumes. Scale bar is 10 μm.

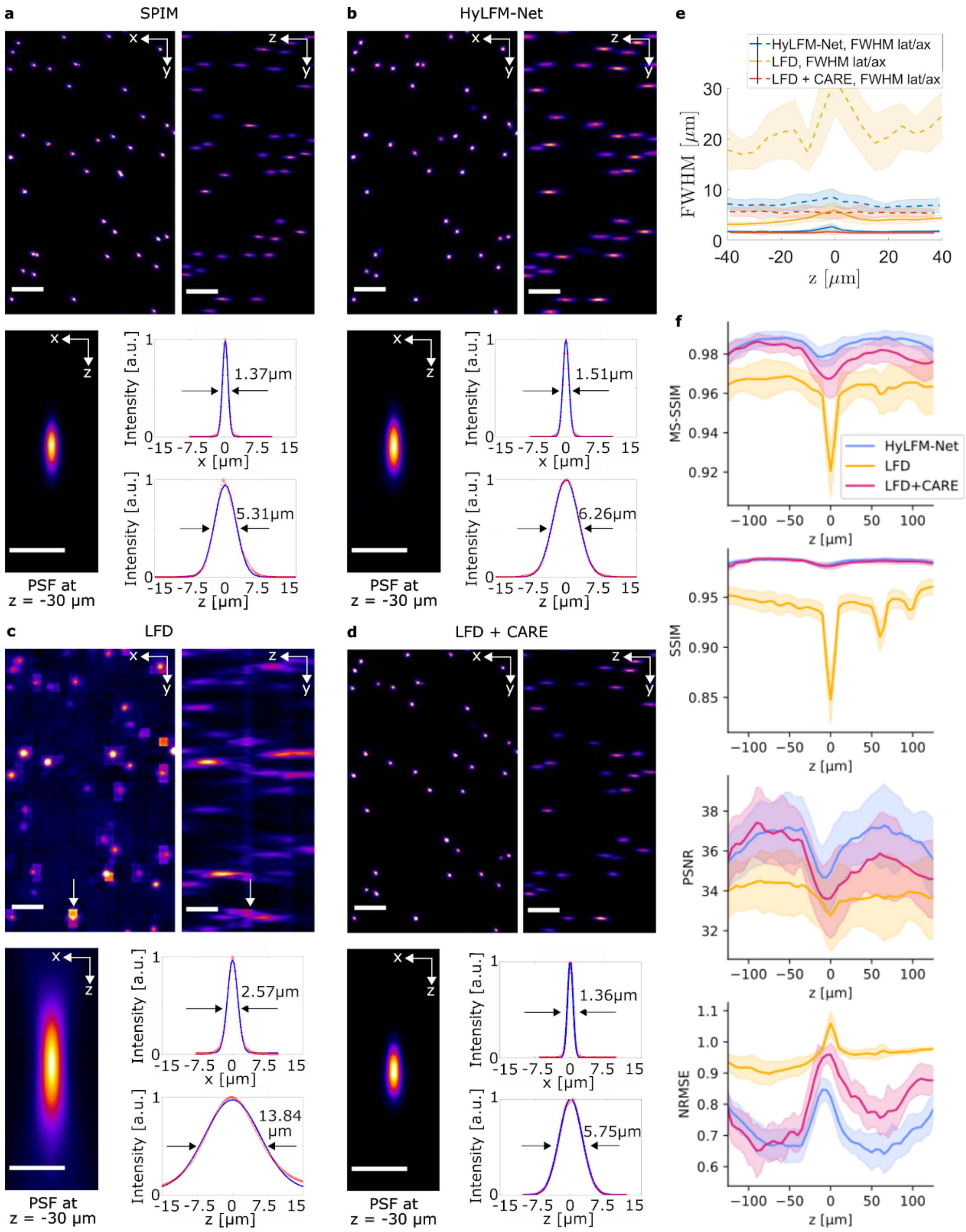


**Supplementary Figure 4: Comparison of HyLFM-Net to a combination of LFD and CARE image restoration on fluorescent beads.** Application of the image restoration CNN CARE (SI Ref.[1], 3D denoising network without any modifications) on LFD-restored volumes improves image quality and metrics with overall performance better than LFD and similar to HyLFM-Net. **(a-d)** Evaluation of SPIM, HyLFM-Net, LFD and LFD+CARE performance on sub-diffraction sized, fluorescent beads, respectively. **(e)** Lateral and axial resolution as a function of imaging depth for SPIM, HyLFM-Net, LFD, and LFD+CARE, respectively. **(f)** MS-SSIM, PSNR and NRMSE image quality metrics across imaging volume comparing the different modalities. Scale bars in **(a-d)** are 20μm in whole FOV (top row) and 10μm in PSF close-up (bottom row).

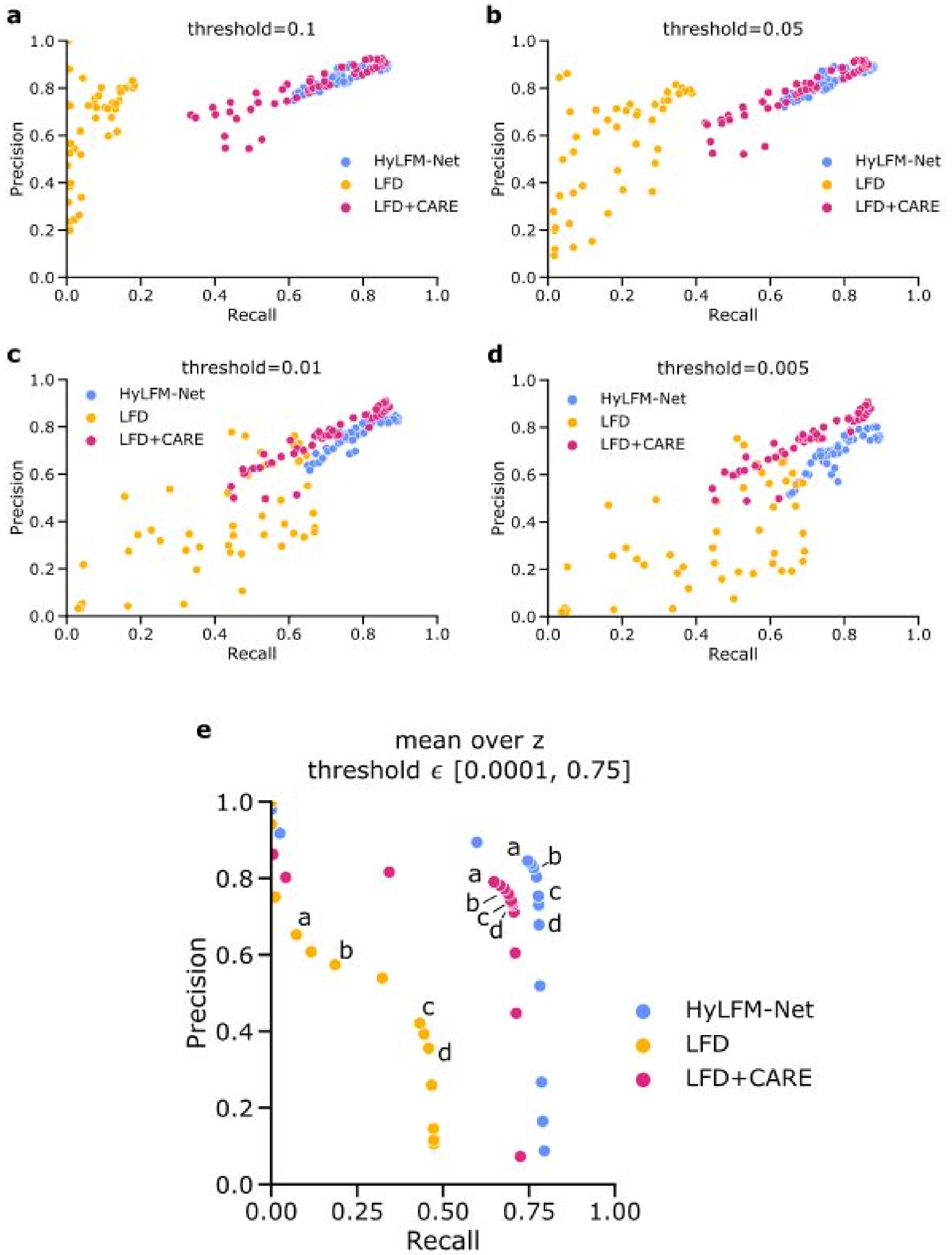


**Supplementary Figure 5: Precision and recall for sub-diffraction beads.**
Precision recall measurements **(a-d)** and curve **(e)** for HyLFM-Net-beads, LFD and LFD+CARE. In **(a-d)** each point represents the average for an individual axial plane. In **(e)** precision and recall were averaged over all volumes, such that each point represents a threshold. All reconstructions were scaled to the SPIM ground-truth to minimize L2 distance; beads were found independently using the Difference of Gaussian (DoG) method with varying thresholds and associated with beads found in SPIM (with threshold 0.1) by Hungarian matching.

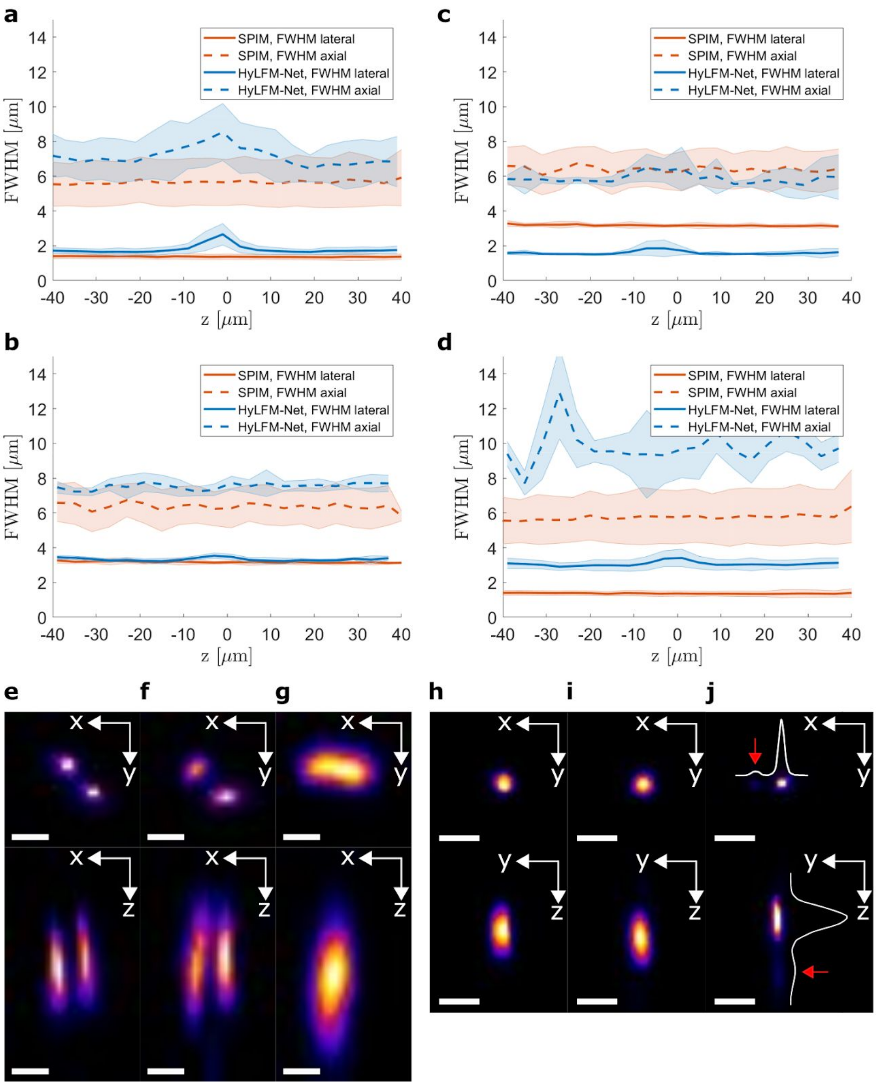


**Supplementary Figure 6: Cross-application of trained deep neural networks can reveal bias to training data.** We created two kinds of samples, one with small (0.1µm) and one with medium-sized (4µm) beads suspended in agarose. In **(a)**, HyLFM-Net was trained on small beads and applied to small beads. FWHM of the beads in the reconstructed volume shows good agreement with SPIM measurements. The same effect is observed in **(b)**, where the network is trained on large beads and applied to large beads. In **(c)**, HyLFM-Net was trained on small beads and used to reconstruct a volume with large beads, resulting in erroneously small bead reconstructions, due to a mismatch between the training and test data. Similarly, in **(d)**, HyLFM-Net trained on large beads and used to reconstruct a volume with small beads produces erroneously large objects. **(e)** SPIM image of 0.1µm beads, **(f)** reconstructions of HyLFM-Net from **(a)**, trained on small beads, **(g)** reconstructions from HyLFM-Net from **(d)**, trained on large beads. **(h)** SPIM image of 4µm beads, **(i)** reconstructions of HyLFM-Net from **(b)**, trained on large beads, **(j)** reconstructions of HyLFM-Net from **(c)**, trained on small beads. Line profile is shown to highlight a reconstruction error (red arrows), where the network reconstructs very small beads (as found in the training data) and produces an additional erroneous peak where none is present in the ground truth SPIM volume. Shadows in **(a-d)** denote standard deviation. Scale bar 2µm in **(e–g)**, and 10µm in **(h–j)**.

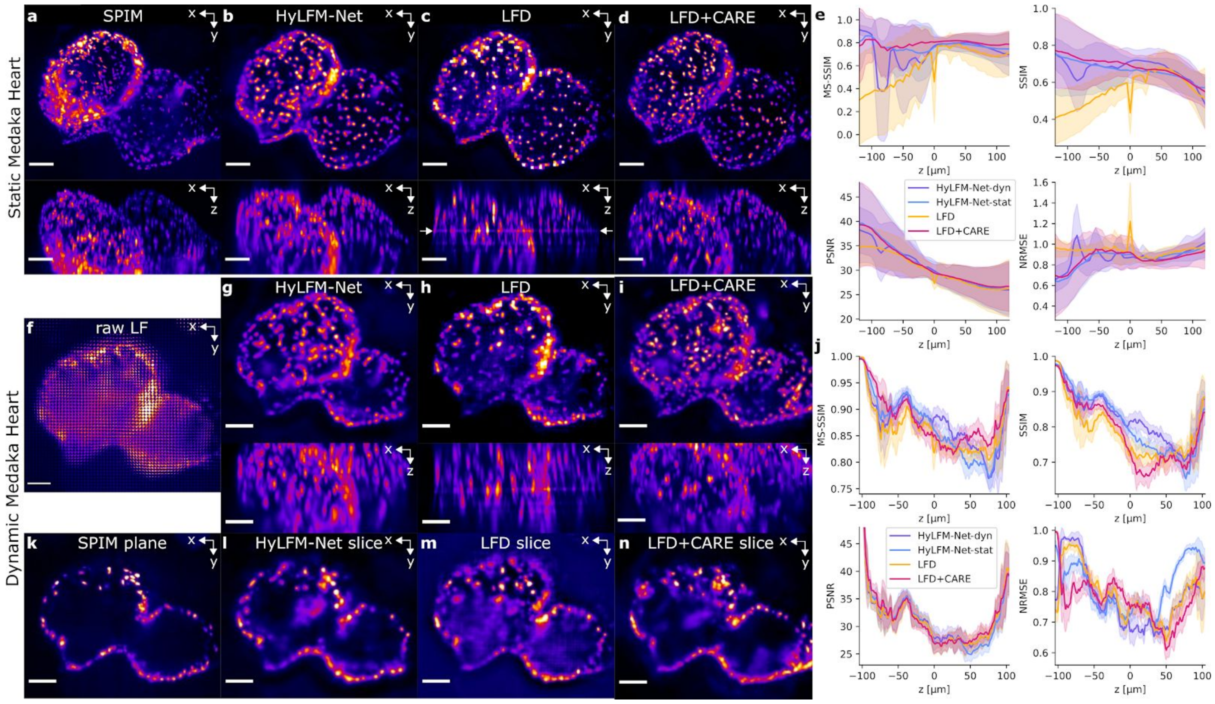


**Supplementary Figure 7: Comparison of HyLFM-Net to LFD and a combination of LFD and CARE image restoration on the heart data.** (**a**) A static hatchling medaka heart acquired by light sheet (SPIM, maximum intensity projections). The corresponding light field volume reconstructions by HyLFM-Net **(b)**, LFD **(c)**, and LFD+CARE image restoration **(d)**. CARE was trained to reconstruct the SPIM volume from the LFD volume. (**e**) MS-SSIM, SSIM, PSNR and NRMSE image quality metrics across the imaging volume of HyLFM, LFD and LFD+CARE restorations compared to SPIM ground truth. **(f)** Example raw light-field (LF) image of a dynamic (beating) medaka heart, acquired at 40Hz. **(g)** Maximum intensity projection of the HyLFM-Net prediction after training on single light sheet planes that continuously swept through the volume during acquisition. Note improved cellular details and absence of artefacts compared to LFD reconstructions in **(h)**. **(i)** Maximum intensity projection of the volume restored by the same CARE network as in **(d)** from LFD reconstructions. **(j)** Image quality metrics MS-SSIM, SSIM, PSNR and NRMSE for single plane data in **(k–n)**. Single plane validation: **(k)** High resolution light sheet image plane at 16μm depth and same time-point as in **(f)** and the corresponding reconstructions by HyLFM-Net **(l),** LFD **(m)** and LFD+CARE **(n)**. HyLFM-Net-stat/-dyn refers to networks trained either on full static volumes of the arrested heart as in **(a–d)** (static medaka heart) or on single planes acquired by imaging a beating heart as in **(k–n)** (dynamic medaka heart). Scale bar is 50μm in **(a–n)**.

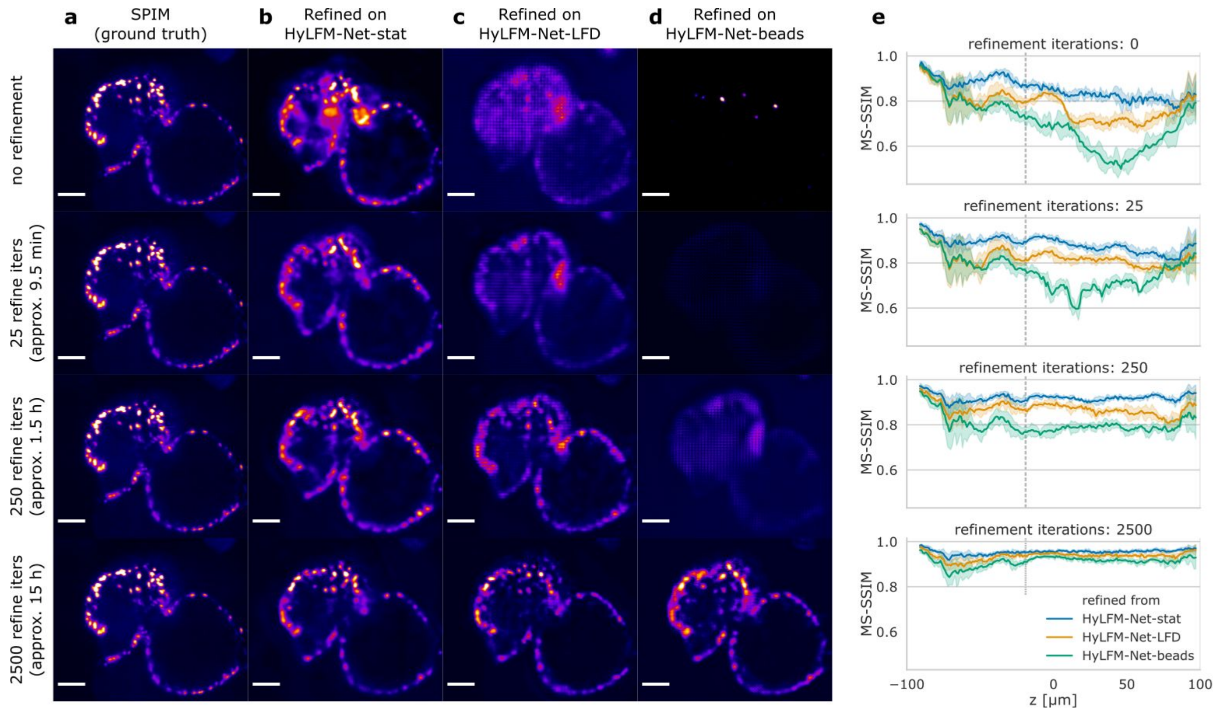


**Supplementary Figure 8: Network fine-tuning for different domain gaps.** Refinement of three differently pre-trained HyLFM networks on dynamically acquired medaka heart images. Column **(a)** shows SPIM ground truth plane at axial position z=-19µm. Columns **(b-d)** depict corresponding slices after increasingly many refinement iterations of different pre-trained HyLFM networks. **(b)** HyLFM-Net-stat (see **Fig. 2b**), **(c)** HyLFM-Net trained on LFD reconstructions of light-field images acquired on another microscope setup (Wagner/Norlin et al, Nat. Meth. 2019 - SI Ref. 2) and **(d)** HyLFM-Net trained on medium-sized beads (see **Fig. 1e**). **(e)** Respective MS-SSIM image quality metrics for each network and stage of refinement. Note that depending on the domain gap, the refinement converges at different speeds, but high fidelity results can be obtained for all pre-trained networks. Also see **Supplementary Video 4**.

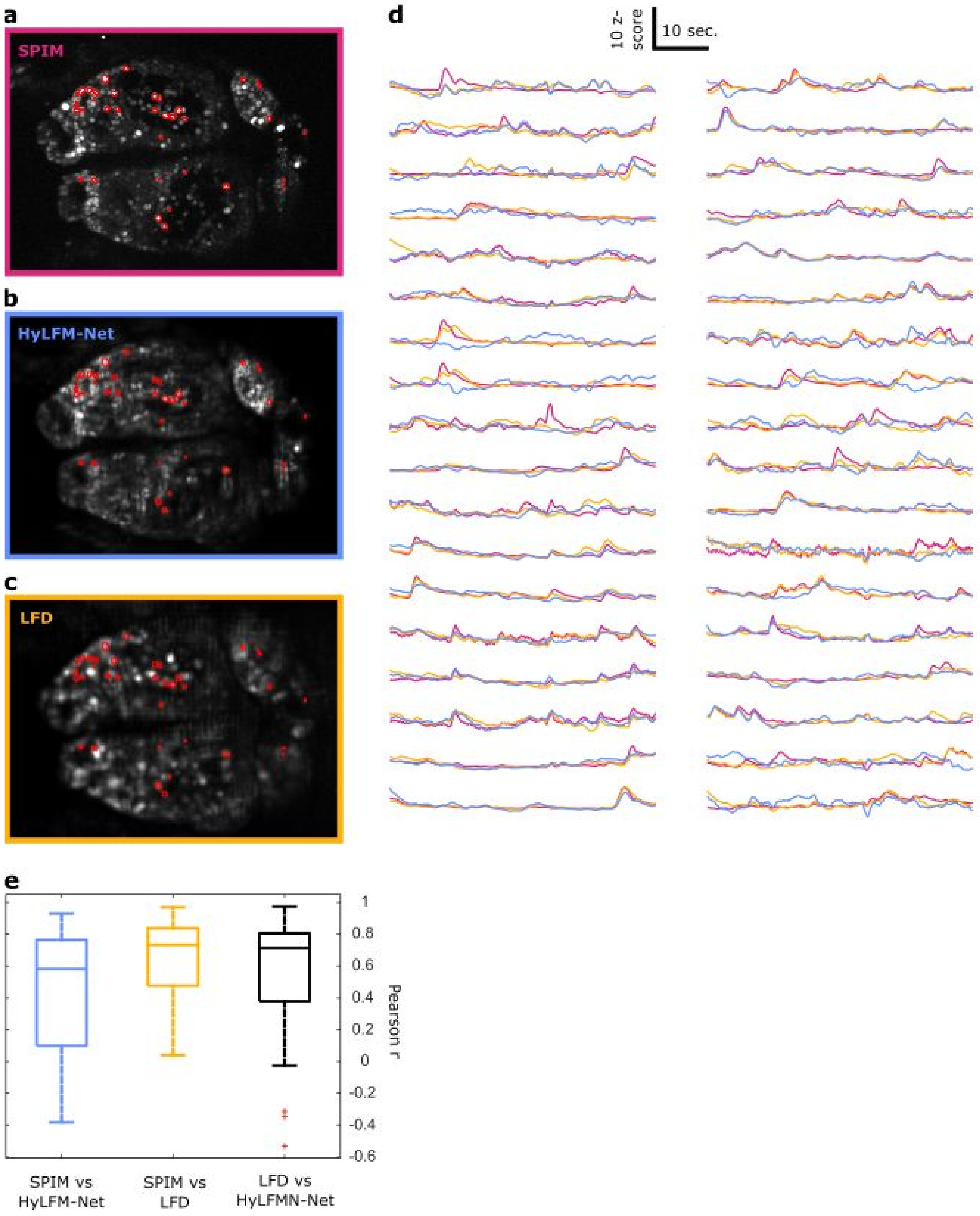


**Supplementary Figure 9: Ca$^{2+}$-imaging in zebrafish larvae brain. (a-c)** Representative image plane acquired with SPIM, HyLFM-Net and LFD, respectively (standard deviation projection over time). **(d)**: Selected Ca$^{2+}$-traces extracted from regions indicated in **(a-c)**. **(e)** Comparison of Pearson correlation coefficients (R) of Ca$^{2+}$-traces extracted by SPIM, HyLFM-Net and LFD. Note that the difference between LFD and HyLFM-Net performance is not statistically significant (p=0.053, Dunn-Sidak). See Methods for details.

| **Medaka heart** | network architecture | | image dimensions | |
|---|---|---|---|---|
| layer name | c | z [px] | y [px] | x [px] |
| input | 361 | 1 | 65 | 75 |
| Res_2d | 488 | 1 | 65 | 75 |
| Res_2d | 488 | 1 | 65 | 75 |
| Up_2d | 244 | 1 | 130 | 150 |
| Res_2d | 244 | 1 | 130 | 150 |
| Proj_2d | 427 | 1 | 130 | 150 |
| Rearrange | 7 | 61 | 130 | 150 |
| Res_3d | 7 | 57 | 126 | 146 |
| Up_3d | 7 | 57 | 252 | 292 |
| Res_3d | 7 | 53 | 248 | 288 |
| Res_3d | 7 | 49 | 244 | 284 |
| Proj_3d | 1 | 49 | 244 | 284 |

| **zebrafish brain** | network architecture | | image dimensions | |
|---|---|---|---|---|
| layer name | c | z [px] | y [px] | x [px] |
| input | 361 | 1 | 70 | 83 |
| Res_2d | 488 | 1 | 70 | 83 |
| Res_2d | 488 | 1 | 70 | 83 |
| Up_2d | 244 | 1 | 140 | 166 |
| Res_2d | 244 | 1 | 140 | 166 |
| Proj_2d | 427 | 1 | 140 | 166 |
| Rearrange | 7 | 61 | 140 | 166 |
| Res_3d | 7 | 57 | 136 | 162 |
| Up_3d | 7 | 57 | 272 | 324 |
| Res_3d | 7 | 53 | 268 | 320 |
| Res_3d | 7 | 49 | 264 | 316 |
| Proj_3d | 1 | 49 | 264 | 316 |

| **beads** | network architecture | | image dimensions | |
|---|---|---|---|---|
| layer name | c | z [px] | y [px] | x [px] |
| input | 361 | 1 | 49 | 74 |
| Res_2d | 976 | 1 | 49 | 74 |
| Res_2d | 976 | 1 | 49 | 74 |
| Up_2d | 488 | 1 | 98 | 148 |
| Res_2d | 488 | 1 | 98 | 148 |
| Up_2d | 244 | 1 | 196 | 296 |
| Res_2d | 244 | 1 | 196 | 296 |
| Proj_2d | 441 | 1 | 196 | 296 |
| Rearrange | 7 | 63 | 196 | 296 |
| Res_3d | 7 | 59 | 192 | 292 |
| Up_3d | 7 | 59 | 384 | 584 |
| Res_3d | 7 | 55 | 380 | 580 |
| Res_3d | 7 | 51 | 376 | 576 |
| Proj_3d | 1 | 51 | 376 | 576 |

**Supplementary Table 1: Network architecture and image dimensions.**
Network layers and their corresponding image/tensor dimensions for medaka heart, zebrafish brain, and beads samples. For medaka heart samples also slightly smaller image dimensions were used (e.g. **SI Video 2,3**). Layer names as indicated in **SI Fig. 1**. For training on single plane images, an additional layer enacting the affine transformation was appended after the final projection.

| Name | LFM image size [px] | Prediction size [px] | Prediction size [µm] | Sample rate [Hz] | Input rate [Mpx/s] | Output rate [Mvx/s] |
|---|---|---|---|---|---|---|
| **HyLFM-Net** | | without i/o to/from hard drive (avg. of 3 runs of 1000 samples) | | | | |
| heart | 1235×1425 | 244×284×49 | 339.0×394.6×245.0 | 18,2 | 32,1 | 61,9 |
| brain | 1292×1577 | 256×316×49 | 355.7×439.1×140.0 | 15,3 | 31,1 | 60,5 |
| beads | 931×1406 | 376×576×51 | 261.2×400.2×100.0 | 5,4 | 7,1 | 59,6 |
| **HyLFM-Net** | | with i/o to/from hard drive (avg. of 3 runs of 100 samples) | | | | |
| heart | 1235x1425 | 244×284×49 | 339.0×394.6×245.0 | 8,2 | 14,4 | 31,3 |
| brain | 1292x1577 | 256×316×49 | 355.7×439.1×140.0 | 9,4 | 19,1 | 41,5 |
| beads | 931x1406 | 376×576×51 | 261.2×400.2×100.0 | 4,3 | 5,7 | 47,7 |
| **LFD** | | without i/o to/from hard drive (avg. of 3 runs of 1 sample with 8 iterations) | | | | |
| heart | 1235×1425 | 1235×1425×49 | 377.9×416.8×245.0 | 9,5E-4 | 1,0E-3 | 5,1E-2 |
| brain | 1292×1577 | 1292×1577×49 | 377.9×461.3×245.0 | 9,0E-4 | 1,1E-3 | 5,3E-2 |
| beads | 931×1406 | 931×1406×49 | 272.3×411.3×245.0 | 9,3E-4 | 1,0E-3 | 5,3E-2 |

**Supplementary Table 2: Inference speed on single GPU**. Speed of HyLFM-Net inference and LFD deconvolution on a Nvidia GeForce RTX 2080 Ti. For LFD we used the following settings: magnification 22.5x, $n_{num}$=19. For HyLFM-Net without i/o and LFD a mini batch size of 1 was chosen. For HyLFM-Net with i/o mini batch sizes of 8, 5, and 1 for heart, brain, and beads were chosen, respectively.

**Supplementary Video 1: Volumetric HyLFM reconstruction of a beating medaka heart at 40Hz.**
Volumetric reconstruction of the medaka heart at 40Hz image acquisition speed shown in **Fig. 2(d–i)** and refined HyLFM-Net-stat. The cyan plane corresponds to the sweeping SPIM image plane. The panels from left to right show the overlay of the light sheet plane with the respective plane from the HyLFM-Net/LFD volume, a projection of the HyLFM-Net/LFD volume rotated by 45 degrees around the y-axis, and a maximum projection of prediction/reconstruction volume along z-, y-, and x-axis, respectively. Scale bar 30 µm.

**Supplementary Video 2: Single plane HyLFM reconstruction of a beating medaka heart at 56Hz.**
Single plane comparison of SPIM ground truth to the corresponding plane of the prediction volume of HyLFM-Net and reconstruction volume of LFD at indicated axial positions of the medaka heart at 56Hz image acquisition speed. See also **Fig. 2i–k**. Scale bar 30 µm.

**Supplementary Video 3: Single plane HyLFM reconstruction of a beating medaka heart at 100Hz.**
Single plane comparison of SPIM ground truth to the corresponding plane of the prediction volume of HyLFM-Net at indicated axial positions of the medaka heart at 100Hz image acquisition speed. Scale bar 30 µm.

**Supplementary Video 4: Demonstration of continuous validation and network refinement.**
Refinement of three differently pre-trained HyLFM networks on dynamically acquired medaka heart images at 40Hz image acquisition speed. Video of refinement experiments in **Supplementary Figure 8** (see figure caption for details).
